# Feature-temporal predictions dynamically modulate performance, feature-based attentional capture, and motor response activity during visual search

**DOI:** 10.1101/2025.04.18.649503

**Authors:** Gwenllian C. Williams, Anna C. Nobre, Sage E.P. Boettcher

## Abstract

Recent research has investigated visual search in dynamic environments and considered how temporal predictions modulate behaviour over time. Spatiotemporal predictions have been shown to adaptively guide behaviour during search to improve target detection at temporally-likely locations. The utility of non-spatial, feature-temporal predictions is less intuitive. The present study investigated whether and how behaviour may be guided toward feature-temporally predictable targets during visual search in crowded and dynamic displays. We tested 1) whether visual attention is guided toward target-relevant features in a temporally specific manner and 2) whether our motor system is temporally tuned accordingly. Participants searched for two distinct targets (non-overlapping colour-shape combinations) in a free-viewing dynamic visual-search task. During an initial learning session, each target identity appeared at a predictable time. In a following testing session, some targets appeared at unpredicted times. Targets’ locations were always unpredictable. We compared the efficiency of identifying targets appearing at predicted versus unpredicted times to test for performance benefits of feature-temporal predictability. In addition, we measured whether gaze fixation was captured by distractors that shared overlapping features with temporally predicted targets. Electromyography recordings were used to test for anticipatory muscle activity tuned to the timing of predictable targets. Results indicated that learning the task regularities led to dynamic modulation of participants’ efficiency at identifying targets, their feature-based attentional capture, and temporally tuned muscle activity. The work highlights the flexibility of temporal expectations in guiding behaviour even under spatial uncertainty.

## Introduction

Visual search is an effective experimental framework for investigating how our attentional system guides the selection of task-relevant stimuli amidst distraction. This guidance relies on the bottom-up saliency of stimuli in the environment, internal representations of target features, and learned environmental regularities (Anderson et al., 2021; Awh et al., 2012; Hollingworth, 2012; Wolfe, 2021; Wolfe & Horowitz, 2017). While most previous visual-search studies have focused on static environments (Wolfe, 2020), real-world settings are often dynamic, with targets and distractors entering our visual field at varying times. More recently, research has incorporated dynamic displays and begun to explore how temporal attention may facilitate target identification within these contexts.

Recent studies have explored how spatial attentional guidance may operate in an adaptive, temporally contingent manner during dynamic visual search. In these studies, participants’ performance benefitted from learning spatiotemporal regularities, where the length of time preceding target appearances predicted their likely spatial locations (Boettcher et al., 2022; Shalev et al., 2022, 2024; Xu et al., 2023, 2025). For example, Boettcher et al., 2022 demonstrated our ability to learn and use spatiotemporal regularities to aid target identification in dynamic and visually competitive environments. Here, participants searched for targets among distractors while stimuli appeared and disappeared transiently at different times and locations during extended trials. Participants were faster and more accurate when identifying targets that consistently appeared at the same time *and* location (e.g., a target that always appeared in the top left, two seconds into a trial) compared to targets that appeared at unpredictable times and locations.

In principle, spatiotemporal regularities may facilitate behaviour through various mechanisms. For example, studies have shown that learning spatiotemporal regularities leads to anticipatory eye movements towards temporally-likely target locations (Boettcher et al., 2022; Pfeuffer et al., 2020). This suggests that we can learn and use spatiotemporal regularities to adaptively focus spatial attention over the course of search. Moreover, some behavioural benefits may result from enhanced response preparation and oculomotor planning. Indeed, temporal expectations are known to act on the motor system, leading to faster responses to temporally expected, compared to unexpected, targets (Rolke & Ulrich, 2010; Thomaschke et al., 2011; Volberg & Thomaschke, 2017).

To better understand the utility of temporal predictions in guiding real-world behaviour, it is essential to consider if temporally-tuned behavioural guidance can also operate effectively under spatial uncertainty. Consider searching for two visually distinct targets in a dynamic scene. It is possible that each target appears with predictable timing but in an unpredictable location. In this case, dynamically shifting attention towards *features* of the temporally-anticipated target, regardless of location, is likely more efficient than continuously attending to features of both targets equally. However, our ability to use such non-spatial, *feature*-temporal regularities to guide visual search behaviour is less well explored than its spatial counterpart. Though few in number, some studies have found initial evidence for attentional guidance based on feature-temporal regularities. Importantly, these studies have used tasks performed at fixation, where stimuli appeared in isolation at just one or two possible locations (Echeverria-Altuna et al., 2024; Thomaschke et al., 2016; Wagener & Hoffmann, 2010). This raises the question of how robust temporal guidance may be under greater spatial uncertainty.

The present study builds on these initial works by investigating feature-temporal guidance in a free-viewing dynamic visual search (DVS) task. Here, target and distractor stimuli could appear concurrently and in locations spanning the display, requiring feature-based attention to operate selectively over a wide spatial range. We investigated whether participants could utilise feature-temporal regularities embedded within the task to guide their behaviour in such a busy dynamic environment filled with unpredictable distraction. This study therefore extends our understanding of whether regularity-based feature-temporal predictions can effectively guide attention in real-world settings where these demands are commonplace. The present study also furthers previous work by recording continuous eye movement and hand muscle activity measures throughout trials, in addition to traditional target-based performance metrics. These additional measures allowed us to more directly examine the dynamic effects of temporal predictions on underlying attentional capture and motor preparation processes during search.

During DVS trials, participants searched through stimuli (coloured shapes) which appeared transiently at different times and locations for two distinct targets among distractors, identifying targets with a key press. Targets had non-overlapping identifying features, while distractors shared zero or one feature with each target. Critically, during the initial learning session, each target appeared predictably at either an early or late moment during the trial. We introduced temporally invalid targets in the subsequent testing session. Independently across all experimental trials, each target stimulus was equally likely to appear, and a target appearance was equally likely to occur early or late. However, taken together, the timing of a target appearance predicted the likely target stimulus.

Using behavioural measures, we first tested whether the feature-temporal regularities influenced participants’ performance in identifying each target. To elucidate the mechanisms underlying behavioural benefits of feature-temporal predictions, we then used eye tracking to test whether gaze would be captured most by distractors that shared an identifying feature with the temporally predicted target. Such a finding would indicate the presence of temporally-tuned feature-based attentional capture. Additionally, electromyography (EMG) data tested whether feature-temporal regularities resulted in anticipatory temporal tuning of motor preparation to respond to the temporally predicted target.

To foreshadow our results, we found that learning feature-temporal regularities dynamically tuned participants’ performance, guided feature-based attentional capture, and modulated anticipatory motor activity. These results provide evidence that humans can utilise non-spatial feature-temporal regularities to guide multiple aspects of behaviour while searching for targets in busy and dynamic environments filled with unpredictable distraction.

## Methods

### Participants

The sample size was selected as a multiple of 12 to meet counter-balancing requirements and was based on a pilot study with similar parameters that revealed a medium-sized effect with 36 participants. Because effect sizes are often overestimated in pilot data, we followed established recommendations to increase the power (Albers & Lakens, 2018; Kumle et al., 2021). Forty-eight participants were tested (35 female, one non-binary, 12 male; mean age = 22.85 ± 4.27 (SD)). We recruited participants aged between 18 and 40 who reported normal or corrected vision (including colour vision), fluency in English, no history of neurological or psychiatric disorder, and not taking any psychoactive medication. All participants gave informed consent. Participants received either financial compensation (£15 per hour) or course credits. All experimental procedures were reviewed and approved by The University of Oxford Medical Sciences Interdivisional Research Ethics Committee.

### Apparatus

Participants sat positioned on a chin rest approximately 95 cm from the monitor (27-inch (68.58 cm) Acer XB270H, refresh rate 100 Hz, 1920 x 1080 pixels resolution). The Eyelink-1000-plus desktop mount (SR Research, Ontario, Canada) was used to record eye movements at 1000 Hz. We used a 9-point calibration routine with an error threshold of .5 degrees of visual angle. Drift correction was applied between each block. When necessary, participants were recalibrated before beginning the next block of trials. The experimental script was generated using MATLAB (2018) and will be made publicly available upon acceptance of the manuscript.

### Dynamic Visual Search Task (DVS)

#### Stimuli

The search display contained a selection of the stimuli, illustrated in Figure 1A. They appeared at various times and locations on a static greyscale 1/F noise background (Figure 1B). A video demonstration of a trial can be found here https://osf.io/bjhf3. Backgrounds were uniquely generated for each trial. This stimulus set consisted of all combinations of three distinct colours (‘pink’ (RGB (255,132,255); ‘blue’ (RGB (1, 230, 207); and ‘orange’ (RGB (255, 147, 0)) and three distinct shapes drawn from equidistant points in the Validated Circular Shape Space (Li et al., 2020). Each stimulus was sized to fill a 54 x 54-pixel box on the display, approximately equal to one degree of visual angle (DVA).

**Figure 1.**
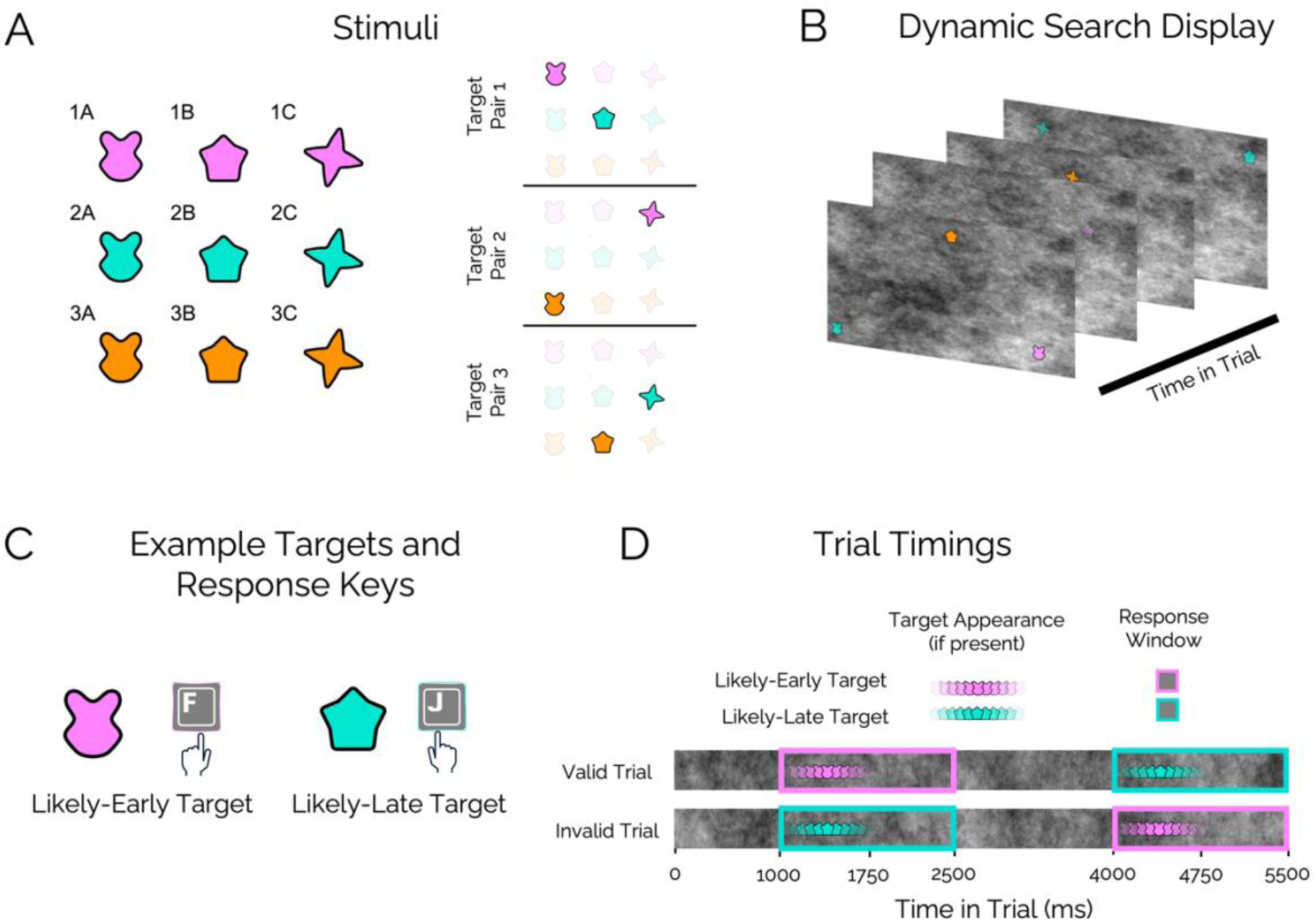
DVS schematic. (A) The nine coloured shape stimuli that appeared as targets or distractors during visual search trials and the three possible target pairs whose assignments were balanced across participants. (B) Illustration of a single DVS trial. The coloured shape stimuli transiently faded in and out of a static noise background at different locations and times during the trials. (C) Example pair of target stimuli, along with their associated response keys and temporal predictabilities. For illustrative purposes, this example is referenced throughout the paper. (D) Schematic of the trial timings for valid trials (top row) and invalid trials (bottom row). Whenever a target appeared, it onset at either 1000 or 4000 ms, and was visible for 750 ms. Participants had 1500 ms from target onset to respond correctly.

Participants were assigned two stimuli as their targets. Targets differed in both colour and shape. The assignment of three possible target pairs was balanced across participants. The pairs were [1A and 2B], [1C and 3A], or [2C and 3B] and are pictured in Figure 1A. The seven non-target stimuli appeared as distractors. Each distractor shared either zero or one feature (colour or shape) with each target. During every trial, exactly 14 stimuli appeared. Zero, one, or two of these stimuli were targets. Each target appeared no more than once in a trial. Each distractor stimulus appeared at least once per trial, and, during trials where no targets appeared, each distractor appeared twice. On trials with one or both targets, six or five distractors, respectively, were randomly selected without replacement to appear a second time.

#### General Timings

All trials lasted 5500 ms. Stimuli faded in from zero to the maximum opacity of 0.75 over 250 ms, remained at maximum opacity for 250 ms, and then faded out to zero opacity over 250 ms. The total duration during which stimulus opacity was greater than zero was 750 ms. ‘Onset time’ of a stimulus is defined as the time the stimulus began fading in relative to the beginning of a trial. Importantly, onset time did not necessarily equate to stimulus visibility.

#### Target Timings (Predictable)

Each of the two possible targets in a trial appeared at a predictable onset time: 1000 ms or 4000 ms. One of the targets, the ‘likely-early target’, was most likely to appear at 1000 ms while the other, the ‘likely-late target’, was most likely to appear at 4000 ms (Figure 1C & 1D). The exact probabilities of the targets’ onset times depended on the experimental session (see ***Learning and Testing Sessions*** below). We did not inform participants of these regularities. The assignment of each target as the likely-early or likely-late target was counterbalanced across participants.

#### Distractor Timings

Distractors were temporally unpredictable. On each trial, the onset time for each distractor was randomly selected, without replacement, from a list ranging from -650 ms to *5*450 ms in 100 ms steps. The negative onset times served as dummy values to allow distractors to be part-way through their appearance at the start of a trial. This assignment resulted in a consistent average level of visual distraction throughout trials. While no two stimuli could onset within 100 ms of each other, multiple stimuli could be visible concurrently given the 750-ms stimulus duration.

#### Stimulus Locations

All target and distractor stimuli were spatially unpredictable across trials. Stimulus locations were pseudo-randomly assigned on each trial. Specifically, locations were assigned such that a stimulus could appear anywhere on the display, provided that none of its edges were within 54 pixels (approximately one DVA) of another stimulus or the edge of the display. The location of each stimulus remained constant as it faded in and out.

#### Identifying Targets

Participants were tasked with finding all targets that appeared during a trial. They were instructed to report targets with a key press as quickly as possible. One target required an ‘F’ response by the left index finger, while the other required a ‘J’ response by the right index finger (e.g., Figure 1C). We counterbalanced the assignment of response keys to each target (likely-early or likely-late) across participants. Participants had a window of 1500 ms following target onset to provide a correct response (Figure 1D). This response window lasted 750 ms after the disappearance of the target to allow for post-stimulus processing and late responses. Any responses made outside of this window were classed as false alarms. Participants were aware that both targets, one target, or no targets could appear on any given trial but were not instructed how many targets to expect on a specific trial.

#### Feedback

After each trial, feedback was provided. “Correct” was displayed whenever correct responses were made to all targets and no false alarms occurred. “Incorrect” was displayed when participants missed a target, responded with the incorrect key, or made a false alarm.

#### Learning and Testing Sessions

Unbeknownst to participants, they completed a learning session followed by a testing session. The learning session consisted of the first four blocks of the DVS, and the testing session consisted of the final block. Each block had 100 search trials randomly intermixed: 25 trials with no targets, 25 with only the likely-early target, 25 with only the likely-late target, and 25 with both targets. Each target, therefore, appeared in an equal number of trials overall.

During learning trials, the time when each target appeared was entirely predictable. If the likely-early target appeared, it always onset at 1000 ms. If the likely-late target appeared, it always onset at 4000 ms. All learning trials were ‘valid’ trials.

During testing, target timings were less predictable. Timings were preserved in 72% of target-present trials (‘valid’ trials) and were reversed in 28% of target-present trials (‘invalid’). During an invalid trial, whenever a particular target appeared, it appeared at the onset time previously associated with the other target. In other words, the likely-early target onset at 4000 ms and the likely-late target at 1000 ms. Figure 1D depicts the trial timings for valid and invalid trials.

### Procedure

Participants first provided informed consent and demographic information before being prepared for EMG recordings (see **EMG Collection**). They then read the DVS instructions and completed a short training session to ensure they knew their targets and response keys. During each of 28 training trials, participants were shown a coloured shape and instructed to respond using the appropriate ‘F’ or ‘J’ key only if that stimulus was a target. Each target was shown seven times and each distractor twice. Participants were required to get 80% of training trials correct to progress, or the session repeated.

Following training, participants completed 16 practice trials of the DVS with valid target timings. These included four trials in which both targets appeared, four with only the likely-early target, four with only the likely-late target, and four target-absent trials, randomly intermixed. After practice, participants completed four learning blocks with 100% valid targets and one testing block with 72% valid targets. Participants could take breaks lasting up to three minutes between blocks. Experimental sessions lasted between 90 and 120 minutes.

### EMG Collection

Muscle activity (EMG) was recorded with SynAmps amplifiers and Neuroscan data acquisition software (Compumedics). Two Ag-AgCl electrodes over the first dorsal interosseus muscle (FDI; between the index finger and thumb) recorded EMG in a bi-polar arrangement from each hand (see Figure 4A). The positive electrode was placed over the muscle belly and the negative electrode over the muscle tendon. A ground electrode was placed above the participant’s left elbow (see Gagné & Schneider, 2008; Kleim et al., 2007; Kwon et al., 2016). Participants were instructed to respond with the left index finger on the ‘F’ key and right index finger on the ‘J’ key to ensure different hands were used to identify the likely-early target and the likely-late target. Electrode impedance was aimed to be below 20 kΩ and was at least below 50 kΩ. Supplementary Figure 1 shows the strong EMG signals recorded during learning trials where two targets appeared, each requiring a response. The relevant analysis focused only on changes in anticipatory EMG muscle excitability during target-absent trials.

### Variables of Interest

Data analysis served three goals. The first question was whether participants learned and used feature-temporal regularities to guide performance. Behavioural data from the testing session most directly addressed this question by testing for performance benefits to identify targets appearing at valid versus invalid onset times. The second question was whether internal templates of the target features guiding search were present and temporally tuned according to the environmental regularities. To address this, we used eye tracking to measure fixations on distractors during target-absent trials in the learning session depending on whether they shared identifying features with the temporally predicted target. Lastly, we investigated whether motor activity related to each target response became temporally tuned to the predictable target timings. The analysis used EMG recordings from both hands in target-absent trials of the learning session.

#### Testing Session: Performance Measures of Feature-Temporal Expectations

The inverse efficiency score metric (IES; Townsend & Ashby, 1983) was chosen as the dependent variable of interest. The IES combines accuracy and RT into one metric. IES was calculated by dividing mean RT by the proportion of target hits. Lower IES thus indicated more efficient performance. Hits were a binary measure indicating whether a target had been correctly identified within the time limit (hit; 1), or not (0). RTs were calculated as the time, in ms, between the onset of a target and the first correct key press which resulted in a hit. As such, a correct classification (hit) was contingent on RT speed. Correct key presses outside the response window (slower than 1500 ms) were not counted as hits. The IES metric, which combines hits and RT, therefore, was especially useful for this study.

Our two main independent variables of interest were target onset (1000 ms or 4000 ms) and target likelihood (likely-early or likely-late). For each participant, we calculated the IES for targets at each combination of levels of these variables (corresponding to each of the bars in Figure 2). Likely-early targets with a 1000 ms onset or likely-late targets with a 4000 ms onset were considered valid. In contrast, likely-early targets with a 4000 ms onset or likely-late targets with a 1000 ms onset were considered invalid. If participants utilised regularity-based predictions, performance should be more efficient when identifying temporally valid versus invalid targets.

**Figure 2.**
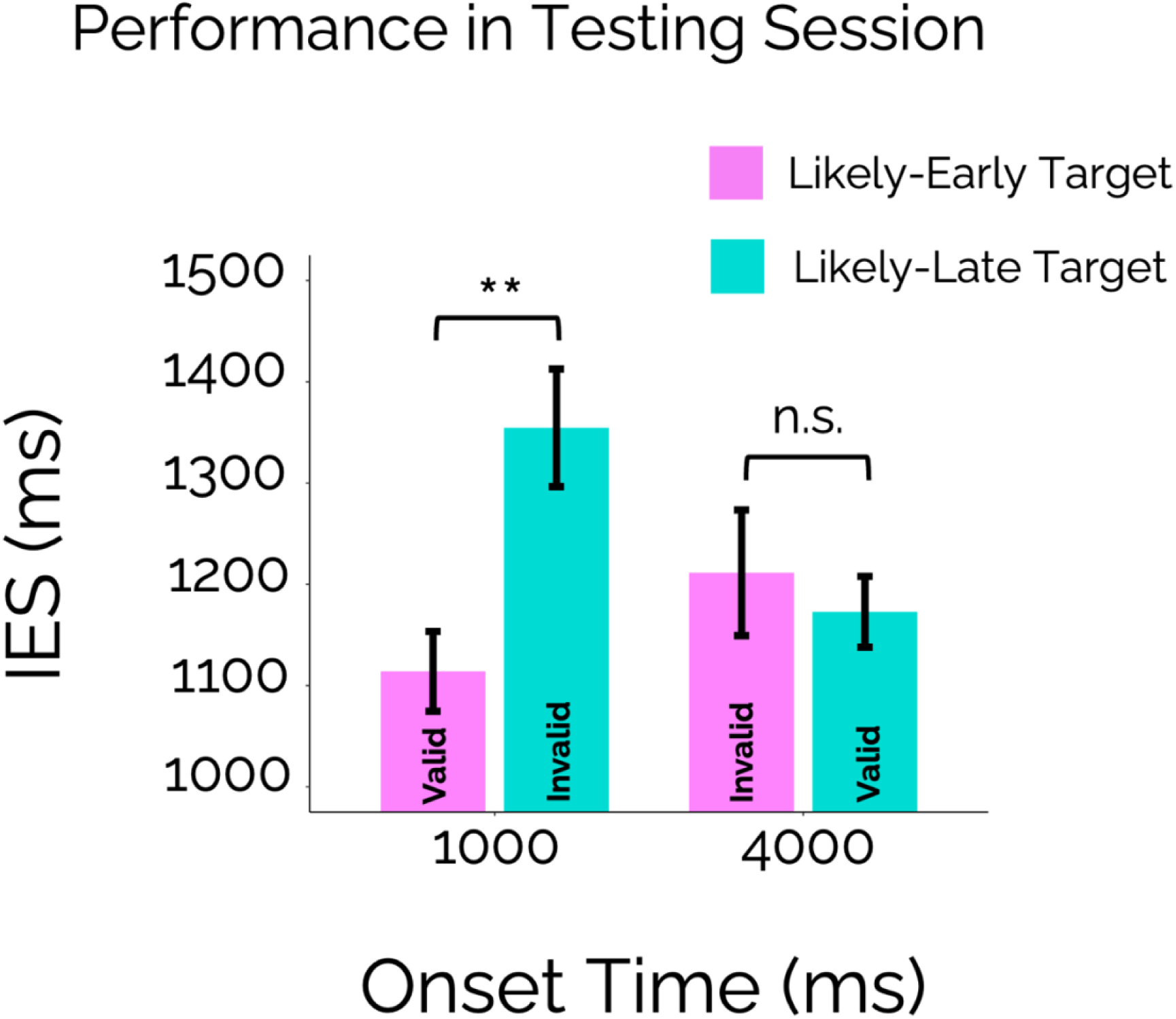
Mean IES for identifying targets in the testing block, for likely-early and likely-late targets appearing at each onset time. Error bars represent ±1 standard error (SE) around the mean. The statistical significance of comparisons is indicated with asterisks: ** = *p* < .01, n.s. = not significant.

We omitted targets that appeared second during two-target trials from the analysis, because the identity of the second target could be considered cued by the identity of the first target. Omitting these targets avoided conflating behavioural effects due to this identity-cueing with effects due to temporal predictions.

#### Learning Session: Feature-Temporal Capture by Distractors

Attentional capture by distractors sharing features with the targets offered a means to test whether participants used regularity-based temporal predictions to tune their visual attentional guidance to relevant features. Specifically, we tested whether eye movements were differentially directed toward the features (colour and shape) of the likely-early target versus the likely-late target at different times throughout trials.

Visual fixations on individual distractor stimuli were the dependent variable that served as a proxy measure of attentional capture. Fixation was a binary measure indicating whether a distractor was fixated (1) or not (0). The premise was that a fixated distractor had captured more attention than a non-fixated distractor (Hollingworth & Bahle, 2020). Fixations were considered as being on a distractor if they occurred within 107 pixels (two DVA) of the stimulus centre and began during the distractor’s 750-ms appearance. Here we used the fixation events and their corresponding start times and average spatial positions recorded by EyeLink. Distractors were classified by whether they shared each feature (colour or shape) with the likely-early, likely-late, or neither target. This resulted in two factors per distractor (colour type and shape type) with three levels each (likely-early target match, likely-late target match, or no match). Figure 3A illustrates the distractors at each level of colour type for our example targets. Additional independent variables were the distractor’s onset time and the learning block (one to four). Note, here we considered the distractor’s onset time – not the absolute time in the trial – to be more directly comparable with how target regularities were defined.

**Figure 3.**
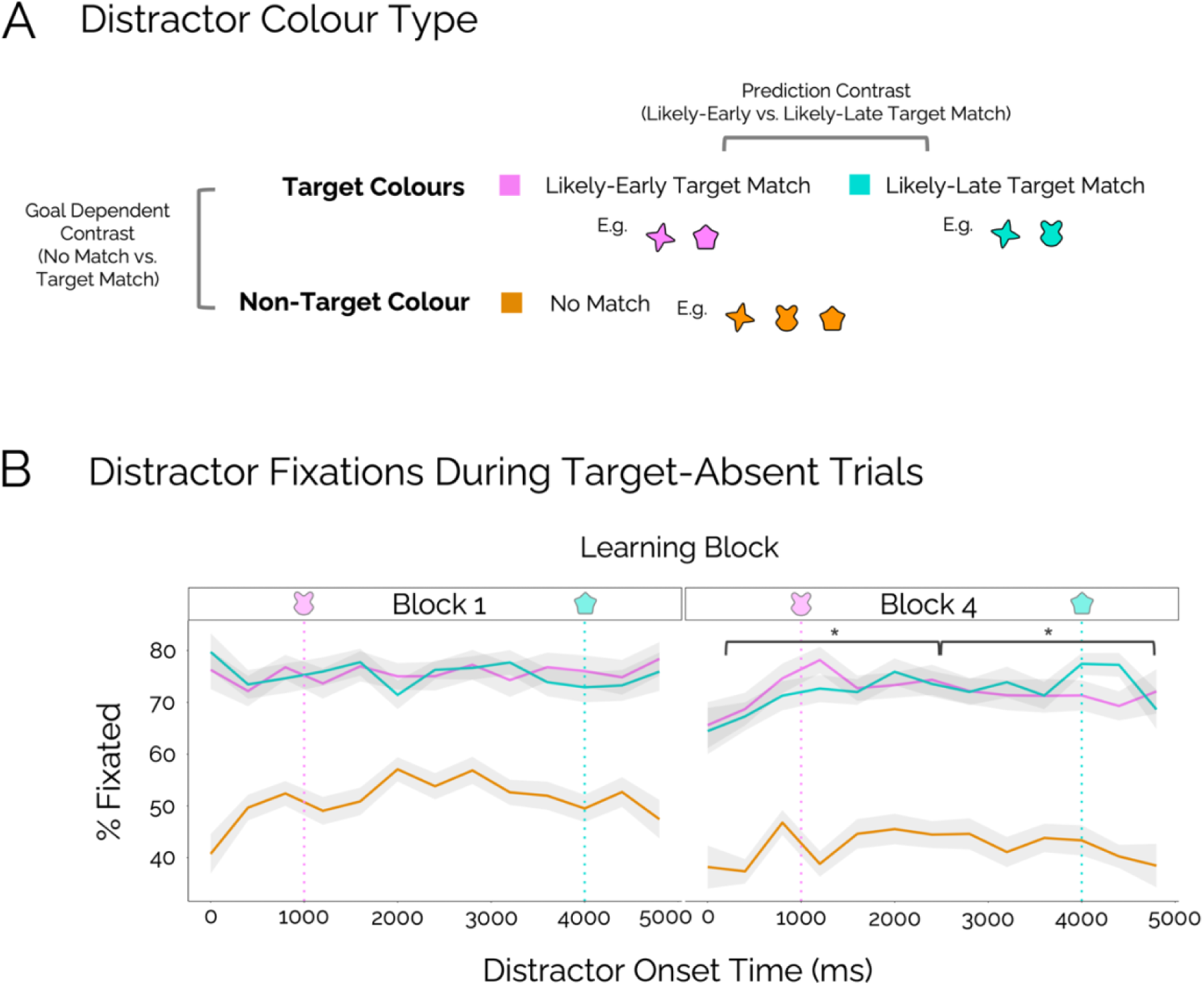
(A) Distractors matching the colour of the likely-early target, the likely-late target, or no target for the example targets from Figure 1C. The two different contrast comparisons for levels of colour type are also illustrated. (B) Percentage of distractors fixated in the first and last learning session blocks, for each level of colour type, across onset times from 0 to 4750 ms. This plot only includes data from target-absent trials. The dotted pink and blue vertical lines indicate the expected onset times for the likely-early and likely-late target, respectively, illustrated using the example targets. The solid pink, blue, and orange lines represent the mean values, while the grey shaded areas represent ±1 SE around the mean. For this visualisation only, data were smoothed by rounding onset times to the nearest 400 ms. The statistical significance of follow-up comparisons between likely-early and likely-late target match distractors in Block 4 is indicated with an asterisk where * = *p* < .05.

A preliminary analysis determined which target features, colour and/or shape, participants used to guide attention during the task. We compared the probability of fixating a distractor which either did or did not share colour or shape with a target. If a particular target feature captured attention, then the distractors with that feature would be more likely to be fixated. Based on this assumption, the preliminary analysis suggested that participants overwhelmingly used target colours, rather than target shapes, as a basis for attentional guidance during the DVS. Therefore, the remainder of the analyses focused on colour type, rather than shape type. Details of this preliminary analysis, and further analyses of shape type, will be made available upon acceptance of the manuscript.

The subsequent, central analysis compared the difference in fixation likelihood between distractors that shared their colour with the likely-early target versus the likely-late target across onset times and learning blocks. We hypothesised that if feature-based attentional guidance followed the task regularities, then distractors that shared their colour with the likely-early target would more likely be fixated when they appeared early in the trial. In turn, distractors that shared their colour with the likely-late target would more likely be fixated when they appeared late. Additionally, the pattern of results should become more pronounced during later learning blocks, as participants gained exposure to the temporal regularities.

Only data from target-absent trials in the learning session were included in these analyses. Target-absent trials were analysed because they provide the most direct measure of how colour-based attentional capture operated across time without possible contamination from changes in strategy or motivation related to finding a target. Only trials from the learning session were analysed to avoid possible changes in temporal predictions following temporally invalid targets (in the testing session). Analysis was further restricted to distractors that appeared for the full 750 ms (i.e., those with an onset time between 0 ms and 4750 ms).

#### Learning Session: Temporally Tuned Response Activity

EMG recordings were used to test whether the predictable timings of targets resulted in changes in the corresponding motor-response preparation. Specifically, we tested whether muscle activity in the response hands varied systematically across time. The dependent variable was the activity in the FDI muscle as recorded from each hand using EMG. The independent variables were the response hand associated with each target (‘target response hand’: likely-early or likely-late) and learning block number (one to four). We additionally considered ‘time in trial’ as measured from the beginning of each trial in ms. Unlike the distractor fixations analysis, in which we investigated changes across *distractor onset times*, the EMG analysis tested for motor-excitability differences across *absolute time in trial*. This is because, unlike our fixation measure, our EMG recordings were not locked to any stimulus appearance.

The analysis tested whether the difference in activity between the two hands was significant at different time points during trials, across the learning blocks. We hypothesised that the tuning of motor preparation according to the task regularities would result in significantly greater activity in the likely-early target response hand near its anticipated onset time of 1000 ms. Similarly, greater activity in the likely-late target response hand would occur around 4000 ms. The pattern should be most pronounced during later learning blocks, after participants had greater opportunity to learn the task regularities.

The analysis was restricted to target-absent trials from the learning session. This avoided conflating any effects of temporal predictions with effects due to target appearances. All EMG signals were analysed, including those that may have corresponded to a false alarm. However, false alarm rates were low, with participants false alarming on 4.29% of target-absent learning session trials on average.

### Statistical Analysis

All analyses used R (R Core Team, 2024) and plots were created using the *ggplot2* package (Wickham, 2016). Full analysis scripts will be made available upon manuscript acceptance.

#### Data Cleaning

Cleaning of behavioural data from the learning and testing sessions followed the same procedure, performed separately. If a participant’s learning session data were excluded, then so were their testing session data, and vice versa. We replaced all data, including eye-tracking and EMG data, from excluded participants and repeated data cleaning until our pre-determined sample size of 48 was reached.

Data from participants with low accuracy were excluded. The accuracy threshold for inclusion consisted of identifying at least 50% of all targets in both the learning and testing sessions. Data were also excluded if the participant’s mean accuracy was below three standard deviations (SDs) of the mean accuracy across all participants in either the learning or testing session. Overall, three participants were excluded for low accuracy. We planned to exclude any participant whose proportion of false alarms was greater than 50% across all key presses, or more than three SDs from the mean proportion across all participants. However, no additional participants were excluded, and proportions were low, with false alarms comprising on average 5.64% of key presses in the final participant sample.

On the target level, data were processed to remove premature or outlier responses. Data with RT < 200 ms were removed. In addition, responses were removed if the RT was slower than three standard deviations from the individual’s overall median RT. These steps removed 0.67% of data from the learning session and 0.71% of data from the testing session. We planned to exclude data from any participant with more than one-third of trials removed, or whose median RT was more than three SDs away from the median RT across all participants in either session. However, this excluded no further participants.

Forty-six of the 48 participants had useable EMG data. All EMG preprocessing was conducted in MNE-Python (Gramfort, 2013). Data were notch filtered at 50 Hz to remove line noise during recording and later high-pass filtered offline at 20 Hz. On the participant level we re-coded data from each hand as coming from the ‘likely-early’ or ‘likely-late’ target response hand, according to whether they used the ‘F’ or ‘J’ key to identify each target. EMG signals were rectified and normalised, within each hand, and within each trial, per participant.

#### Testing Session: Behavioural Performance

We analysed the IES values using a 2x2 (onset time x target likelihood) repeated-measures analysis of variance (ANOVA) test, using the *ezANOVA* function from the *ez* package (Lawrence, 2016). In all cases, a two-tailed 5% error criterion was used to test for significance. We conducted further pre-planned *t*-tests separately analysing the effect of target likelihood at each onset time following any significant interactions between the two variables. These comparisons were supplemented by paired Bayesian *t*-tests on the same IES values using the *BayesFactor* package (Morey & Rouder, 2022). We used the package’s default prior (r = 707) to test the null hypothesis of no difference between the two sets of values against the alternative hypothesis that there is a difference. The Bayes factor (BF_10_) values indicated anecdotal (BF_10_ > 1 and BF_10_ < 3), moderate (BF_10_ > 3 and BF_10_ < 10) or strong (BF_10_ > 10) evidence in favour of the alternative hypothesis, or anecdotal (BF_10_ > 0.33 and BF_10_ < 1), moderate (BF_10_ > 0.1 and BF_10_ < 0.33) or strong (BF_10_ <0.1) evidence in favour of the null hypothesis.

For completion, supplementary analyses assessed the effects of target likelihood and onset time on hits and RT individually. Results of these analyses are visualised in Supplementary Figure 2 and provided in Supplementary Tables 1 and 2.

#### Learning Session: Distractor Fixations

A generalized linear mixed-effects model (GLMM) with binomial distribution tested for effects on distractor fixations. Mixed-effects models can model the contributions of both fixed effects, related to the independent variables, and random effects, related to differences between participants, to variation in the dependent variable (Harrison et al., 2018). Mixed-effects models are also believed to be robust against unbalanced conditions (Kliegl et al., 2010), as are present in our design. We ran our GLMMs using the *glmer* function of the *lme4* package in R (Bates, Mächler, et al., 2015). The model contained fixed effects for colour type, onset time, learning block number, and their interactions.

Onset time and block number entered the model as centred and scaled continuous predictors. We applied custom contrasts to the colour type factor, details of which are given in Table 3 of the Supplementary Materials. These contrasts allowed for two separate comparisons of the three levels of colour type, which are illustrated in Figure 3A. The first contrast, the ‘goal dependent contrast’, compared fixation rates between distractors that *did not* versus *did* share their colour with any target (no colour match vs. target colour match). This comparison determined whether distraction fixations provided a sensitive measure of goal-dependent feature templates. Following this confirmatory analysis, the main comparison of interest was between distractors that shared their colour with the *likely-early* versus the *likely-late* target (likely-early vs. likely-late target colour match). This comparison was specified by the second contrast, the ‘prediction contrast’.

We selected the random-effects structure of the optimal GLMM in line with suggestions by Bates, Kliegl, et al., (2015). The random-effects structure of the initial, full model included intercepts for all participants and by-participant slopes for the effects of colour type, onset time, block number, and all interactions. We tested this full model for over-parameterization using a principal component analysis (PCA) of the random-effects variance-covariance estimates. Here we employed the *rePCA* function of the *lme4* package (Bates, Mächler, et al., 2015). If the model was over-parameterised, we reduced it by removing the random effects component that explained the least variance and did not contribute significantly to the model goodness-of-fit, as indicated by a likelihood ratio test. This process was repeated until we found the optimal model (as reported in the results), for which all random-effects components were supported by the PCA and contributed significantly to the goodness-of-fit.

The results section reports regression coefficients, *β*, alongside the *z*-statistic, and the standard error (SE). The *p*-values are based on asymptotic Wald tests. We used a 5% error criterion to test for significance. Additional pre-planned mixed models helped clarify any significant interactions between the prediction contrast, onset time, and/or block number. These models separately analysed the effect of the prediction contrast for ‘early’ and ‘late’ onset times, and/or for the first and final learning blocks. We define early and late distractor onset times relative to the midpoint between the two target onset times. Early onset times occur before this midpoint (250 – 2500 ms), while late onset times occur after (2500– 4750 ms).

#### Learning Session: Response Hand Activity

Each participant’s EMG data were averaged and then smoothed separately for each learning block. Smoothing was carried out using the *gaussian_filter1d* function from the SciPy library (Virtanen et al., 2020) across a 70-ms window. We tested for significant differences in activity between each target response hand, across time in trial, and across learning blocks by employing the *clusterlm* function from the *permuco* package in R (Frossard & Renaud, 2021). We used a significance level of 5% and ran 5000 permutations. Block entered the model as a continuous variable, centred and scaled. The interaction between block number and the difference between target response hands was also included in the model.

## Results

### Testing Session: Performance Benefits for Feature-Temporally Predictable Targets

The ANOVA revealed a significant two-way interaction between target likelihood and onset time (*p* = .002; Figure 2). All other predictors were non-significant. Table 1 contains the full ANOVA output.

**Table 1.** Testing Session IES ANOVA Output.

| Predictor | $F$ | $df_1$ | $df_2$ | $p$ | $\eta^2_G$ |
| --- | --- | --- | --- | --- | --- |
| Onset Time | 1.08 | 1 | 47 | .304 | .003 |
| Target Likelihood | 2.57 | 1 | 47 | .116 | .019 |
| **Onset Time : Target Likelihood** | **10.28** | **1** | **47** | **.002** | **.035** |

*Note.* Significant predictors are shown in bold.

Two follow-up repeated-measures *t*-tests separately analysed the effect of target likelihood at each onset time. When targets appeared at 1000 ms, target likelihood significantly affected IES, *t*(47) = 3.17, *p* = .003, *d* = 0.46, *BF_10_* = 12.07. As hypothesised, participants were more efficient at identifying the likely-early target (*M_IES_* = 1114 ms, *SD_IES_* = 371 ms) than the likely-late target (*M_IES_* = 1355 ms, *SD_IES_* = 371 ms). In contrast, when targets appeared at 4000 ms, target likelihood had no significant effect on IES, *t*(47) = 0.50, *p* = .620, *d* = 0.07, *BF_10_* = 0.18.

### Learning Session: Fixation Likelihood Across Distractor Colours

When comparing fixation likelihood for distractors according to the colour they shared with targets, the optimal GLMM had a random-effects structure containing intercepts for individual participants and by-participant slopes for colour type (with both goal dependent and prediction contrasts), block number, and their interaction. This model was specified in the syntax of R as: Fixation ∼ Colour Type * Onset Time * Block Number + (1 + Colour Type * Block Number | Participant)

The fixed effects output of this model is given in Table 2 below and visualised in Figure 3 of the Supplementary Materials.

**Table 2.** Optimal GLMM Fixed Effects Output for Distractor Fixation Likelihood.

| Predictor | $\beta$ | $SE$ | $z$ | $p$ |
| --- | --- | --- | --- | --- |
| **Goal Dependent Contrast (No Colour Match vs. Target Colour Match)** | **-0.86** | **0.05** | **-17.26** | **< .001** |
| Prediction Contrast (Likely-Early vs. Likely-Late Target Colour Match) | -0.03 | 0.07 | -0.51 | .608 |
| **Onset Time** | **0.03** | **0.01** | **2.97** | **.003** |
| **Block Number** | **-0.09** | **0.03** | **-3.36** | **< .001** |
| Goal Dependent Contrast (No Colour Match vs. Target Colour Match) : Onset Time | < -0.01 | 0.01 | -0.03 | .973 |
| **Prediction Contrast (Likely-Early vs. Likely-Late Target Colour Match) : Onset Time** | **.07** | **0.03** | **2.50** | **.012** |
| **Goal Dependent Contrast (No Colour Match vs. Target Colour Match) : Block Number** | **-0.05** | **0.02** | **-3.03** | **.002** |
| Prediction Contrast (Likely-Early vs. Likely-Late Target Colour Match) : Block Number | -0.01 | 0.04 | -0.14 | .887 |
| Onset Time : Block Number | 0.01 | 0.01 | 0.99 | .322 |
| Goal Dependent Contrast (No Colour Match vs. Target Colour Match) : Onset Time : Block Number | -0.01 | 0.01 | -1.10 | .270 |
| **Prediction Contrast (Likely-Early vs. Likely-Late Target Colour Match) : Onset Time : Block Number** | **0.08** | **0.03** | **2.98** | **.003** |

#### Goal-Dependent Capture

The GLMM provided evidence for goal-dependent capture. Overall, the likelihood of fixating a distractor was significantly predicted by whether it shared its colour with a target (*p* < .001). Consistent with attentional guidance toward target colours, participants fixated significantly more distractors of target colour (*M* = 72.72%, *SD* = 7.53%) than of non-target colour (*M* = 46.09%, *SD* = 7.53%; Figure 3).

The two-way interaction between the goal dependent contrast and block number was also significant (*p* = .002). The difference between the fixation rate for target colour match vs. no colour match distractors was 23.62% in the first learning block, compared to 30.02% in the final block. This is consistent with participants avoiding fixating more *no match distractors* as the learning session progressed.

#### Feature-Temporal Capture

The GLMM analysis further provided evidence for temporally tuned capture according to the colour of the target predicted at early vs late onset times. The overall likelihood of fixating a distractor was not significantly predicted by whether it shared its colour with the likely-late or likely-early target (*p* = .608). However, importantly, the two-way interaction between the prediction contrast (likely-early vs. likely-late target colour match) and onset time was significant (*p* = .012), indicating that fixations on distractors of each target colour varied depending on their onset time. A further, significant three-way interaction revealed that this two-way interaction between the prediction contrast and onset time depended significantly on block number (*p* = .003).

To interpret the significant three-way interaction, two additional GLMMs analysed the interaction between the colour type prediction contrast and onset time during the first and last learning blocks individually. Both optimal GLMMs had random-effects structures containing intercepts for individual participants and by-participant slopes for colour type:

Fixation ∼ Colour Type * Onset Time + (1 + Colour Type | Participant)

In Block 1, the interaction between the prediction contrast and onset time was not significant (*β* = -0.08, *SE* = 0.06, *z* = -1.36, *p* = .174; Figure 3B). In contrast, and in line with our hypotheses, in Block 4, this interaction was significant (*β* = 0.15, *SE* = 0.06, *z* = 2.71, *p* = .007; Figure 3B). To interpret this significant two-way interaction, we ran two further GLMMs analysing the effect of the prediction contrast during Block 4, for distractors with either ‘early’ or ‘late’ onset times. Early onset times were between 250 and 2500 ms, and late onset times were between 2500 and 4750 ms. Both optimal GLMMs had random-effects structures containing intercepts for individual participants and by-participant slopes for colour type:

Fixation ∼ Colour Type + (1 + Colour Type | Participant)

In learning Block 4, distractors that appeared early during a trial were significantly more likely to be fixated if their coloured matched the likely-early target (*M* = 74.28%, *SD* = 10.00%) rather than the likely-late target (*M* = 70.74%, *SD* = 8.20%; *β* = -0.24, *SE* = 0.10, *z* = -2.37, *p* = .018). In contrast, for distractors that appeared late, those sharing the likely-late target colour (*M* = 73.75%, *SD* = 8.74%) were significantly more likely to be fixated than those sharing the likely-early target colour (*M* = 70.93%, *SD* = 9.26%; *β* = 0.21, *SE* = 0.10, *z* = 2.04, *p* = .042). Overall, by the final learning block, participants demonstrated a preference to fixate likely-early target colour distractors at the beginning of trials and likely-late target colour distractors toward the end of trials (Figure 3B, Block 4).

### Learning Session: Temporally Tuned Response Hand Activity

EMG recordings showed temporally specific modulation of motor response activity for targets predicted to appear at the early interval. Regarding the difference in activity between target response hands, our analysis revealed a significant cluster for which the activity difference peaked at 1466 ms (*p* = .027). Here, participants had significantly greater activity in their likely-early target hand than their likely-late target hand, with the peak difference aligning temporally with the anticipated likely-early target appearance (Figure 4B). A second cluster, corresponding to a later time in the trial was also revealed (time of peak difference 4721 ms). Here, the average activity in the likely-late target hand was greater than that in the likely-early target hand, though this difference was non-significant (*p* = .578).

**Figure 4.**
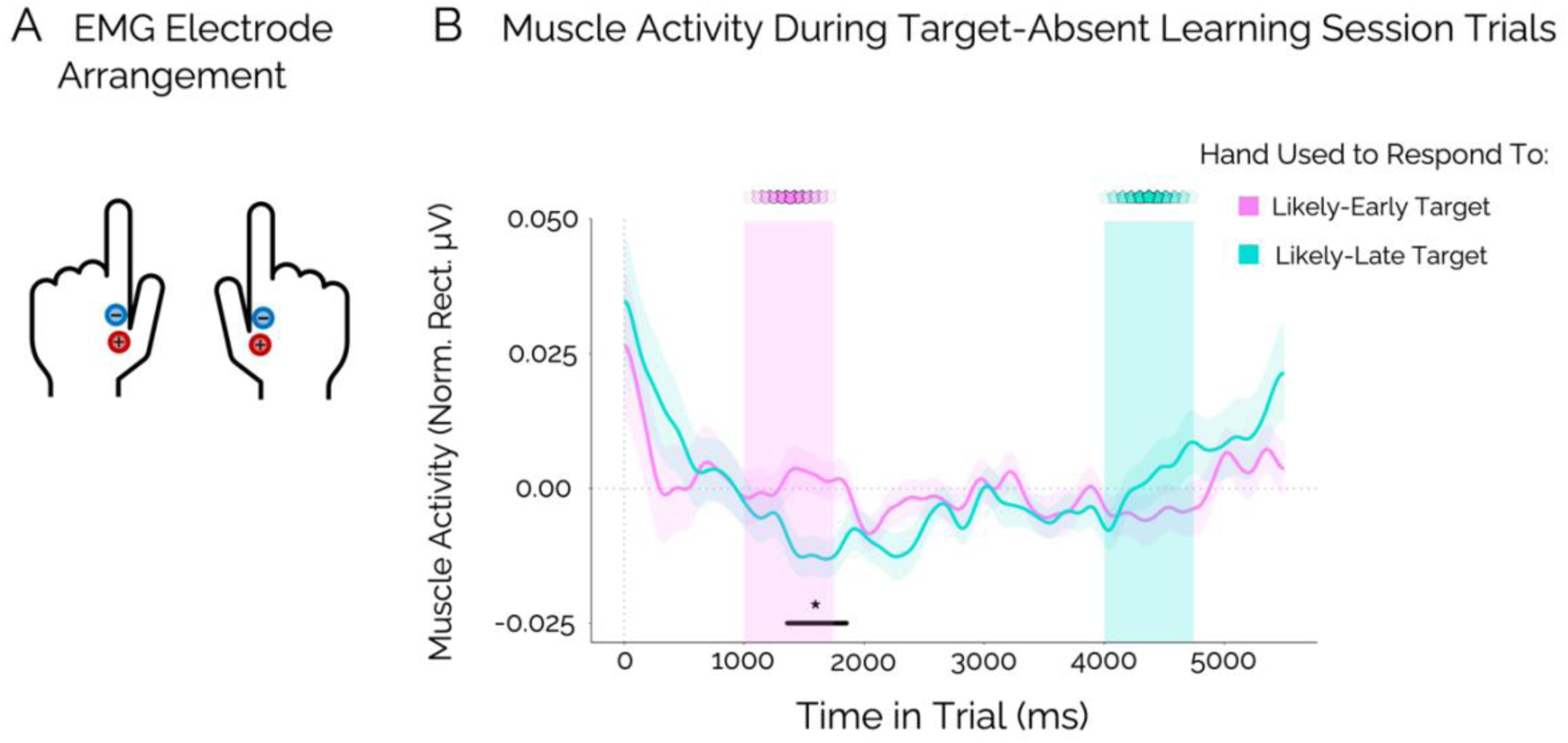
(A) Illustration of the arrangement of positive and negative EMG electrodes over the FDI muscle in each hand. (B) Normalised rectified muscle activity in hands used to respond to the likely-early and likely-late targets over the 5500 ms duration of target-absent trials during the learning session. The solid pink and blue lines represent the mean values and the shaded areas around them represent ±1 SE. The pink and blue columns indicate the time windows over which the likely-early and likely-late targets, respectively, would have faded in and out. The black horizontal line marks a significant cluster where * = *p* < .05.

No significant clusters were found as a function of block number or the interaction between block number and target response hand.

### General Discussion

The present study demonstrates that regularity-based feature-temporal predictions can adaptively guide visual search behaviour across time. Converging behavioural, eye-tracking, and EMG data revealed temporal tuning of participants’ task performance, feature-based attentional capture, and motor activity, in line with task regularities. Extending prior research on dynamic visual search (Boettcher et al., 2022; Shalev et al., 2022, 2024; Xu et al., 2023), our findings highlight the flexibility of temporal attention to modulate search behaviour even under spatial uncertainty. Moreover, our novel paradigm advances previous work on feature-temporal attention (Echeverria-Altuna et al., 2024; Thomaschke et al., 2016; Wagener & Hoffmann, 2010) by demonstrating selective feature-temporal attentional guidance impacting sensory selection and motor preparation to identify and respond to targets appearing among competing distraction in a free-viewing dynamic visual search task.

Our analysis of performance during the testing session revealed that target identification efficiency depended on whether targets appeared at their predicted (valid) or unpredicted (invalid) time. Performance was higher when targets upheld, as opposed to violated, the feature-temporal regularities from the preceding learning session. This finding suggests participants learned and used the task regularities to guide their behaviour. As we only analysed one-target trials, the observed differences in performance for each target between early and late onset times cannot not be attributed to a change in strategy following the detection of a target at the early time. Instead, these shifts are likely the result of participants tracking time within the trial and shifting their internal templates accordingly.

Interestingly, we found that performance differences between feature-temporally valid and invalid targets were most pronounced when targets appeared early. This aligns with many previous studies investigating temporal expectations which found a significantly larger effect following a short, but not long interval (Coull et al., 2000; Heideman et al., 2018; Xu et al., 2023, 2025). Future research should explore why temporal expectations may guide behaviour more effectively at shorter intervals. Factors to consider may include differences in estimating short versus long intervals, ceiling effects at later times, or difficulties enacting shifts in attentional or motor processes away from initial prioritisations.

In contrast, Boettcher et al. (2022) and Echeverria-Altuna et al., (2024) reported significant performance benefits for temporally-expected target locations and features, respectively, occurring at both early and late times during trials. A possible explanation for these differences is that, in the present study, we analysed performance for late targets that were not preceded by another target. For Boettcher et al., 2022, and Echeverria-Altuna et al., (2024), temporally expected late targets were always, or mostly, preceded by an earlier temporally expected target in the same trial. It is possible that these extra temporally predictable targets helped participants to encode temporal regularities at later times by providing additional reliable temporal reference points after trials had begun, and by creating more distinct representations of the early vs. late target timings (Bangert et al., 2020; Ongchoco et al., 2023). Further investigation is needed to explore why effects of temporal predictions may be weighted toward early time points in some contexts and not others, and whether the formation of temporal predictions at later intervals is hindered in environments with fewer reliable temporal anchors.

Going beyond traditional performance metrics, we used eye tracking to probe feature-based capture of attention throughout trials, even in the absence of target appearances. Participants were overall more likely to fixate distractors that shared their colour with one of the targets compared to distractors that did not share their colour with either target. This was an important proof of concept and provided initial evidence that our measure of distractor fixations could reliably track the participants’ internal target template. Without considering onset time, we found no significant difference in the fixation likelihood of distractors which shared their colour with the likely-early target and the likely-late target, suggesting both target colours captured attention to a similar extent overall. However, when considering distractor onset time, we found that, by the final learning block, participants’ attentional capture had become dynamically tuned towards distractors with the same colour as the temporally anticipated target. Importantly, since this analysis focussed on target-absent trials, this effect could not be due to participants having already found a target. Instead, our results suggest dynamic feature-based attentional guidance based on the time in trial. As such, the performance benefits for temporally valid targets during the testing session may have (partially) resulted from greater attentional guidance towards temporally expected target colours.

Our eye-tracking results complement previous work demonstrating anticipatory eye movements toward temporally likely target locations, rather than features, following learning of spatiotemporal regularities (Boettcher et al., 2022; Pfeuffer et al., 2020). One mechanism by which spatiotemporal regularities may benefit behaviour is temporally tuned ocular motor planning. It is not possible to determine the extent to which the results of the aforementioned studies reflect underlying shifts in spatial attention or saccade preparation. In contrast, the present study investigated eye movements in a context where all stimuli appeared with large spatial uncertainty. As such, our findings are unlikely to reflect the temporal preparation of directional saccades. Instead, the gaze results expose more directly dynamically tuned shifts in feature-based attention.

These findings of temporal expectations modulating selective attentional guidance under such large spatial uncertainty extend previous research on temporal attention. They demonstrate that temporal predictions serve to modulate not only where, but also *what* receives attentional priority. Modulation by temporal expectation occurs even when spatial locations are highly unpredictable, beyond just one or two possible locations (Echeverria-Altuna et al., 2024; Thomaschke et al., 2016; Wagener & Hoffmann, 2010). This furthers the argument that, while spatial certainty may aid the operation of temporal attention, it is not always necessary.

While all of our measures revealed behavioural shifts according to the task regularities, our eye-tracking results differed from our performance and EMG findings in one respect. Gaze modulation pointed to significantly greater attentional capture for temporally predicted target colours for early times during trials *and* a (marginally) significantly greater prioritisation of the likely-late target colour over the likely-early target colour was also found for late times during trials. Therefore, while we did not find temporally tuned behavioural benefits, or differences in muscle activity at the late time point, we did find these differences in the eye-tracking results. Further research may explore whether the effects of temporal predictions on earlier stages of target selection, as indexed by the gaze effects, differ in nature from those at later, motor-related processes.

Beyond our primary focus on effects of feature-temporal expectations, our eye tracking analysis revealed a striking effect of task goals on attentional capture. Participants overwhelmingly fixated more distractors that shared their colour with one of the targets, regardless of timing, and this preference increased as they gained task experience. As stimuli appeared across a wide range of unpredictable spatial locations, this supports evidence that goal-driven feature-based attention operates effectively across spatially unattended locations (Brignani et al., 2010; Saenz et al., 2002; Treue & Trujillo, 1999). Additionally, we observed these strong feature-based contingent capture effects in an extended dynamic setting where all stimuli appeared at different times. It is not necessarily intuitive that we should have detected these effects over and above any capture effects due to abrupt stimulus onsets, which have been demonstrated to exist regardless of stimulus task-relevance (Schreij et al., 2008, 2010).

While our study primarily examined colour-based attentional guidance, future research could explore visual attentional guidance toward other features in dynamic contexts, and whether they may be temporally tuned. We found that target shapes did not substantially guide attention in our task, nor was guidance toward shapes temporally tuned (a link to this additional analysis will be made available upon manuscript acceptance). This is perhaps unsurprising, given the dominance of colour in guiding attention compared to shapes (Alexander et al., 2019; Wolfe & Horowitz, 2017). However, it is possible that shape-based guidance may become temporally tuned in a task where shape is the primary basis for guidance. DVS tasks can also provide insights into attentional guidance to dynamic stimulus features, such as motion (Boettcher & Nobre, 2025), or systematically changing colours (Muhl-Richardson, Cornes, et al., 2018; Muhl-Richardson, Godwin, et al., 2018). While our current findings suggest that temporal predictions can shift attentional priorities between internal representations of static target colours, it would be interesting to consider if the same is true for representations of dynamic features.

In addition to performance and eye tracking metrics, our EMG results are consistent with temporal tuning of participants’ behaviour according to task regularities. Our EMG results from the learning session in target-absent trials showed greater muscle activity in the response hand for the temporally expected target, relative to the temporally unexpected target. These findings compliment Volberg & Thomaschke’s (2017) findings of temporally-tuned effector-specific preparatory brain activity in a task where target stimuli and their associated responses varied predictably with the duration preceding the target. Our effect was significant only at early times during trials. Further, this finding did not change significantly across learning session blocks, suggesting it may have been quickly established.

Within our task, specific motor responses were inherently tied to the feature-temporal regularities, which may have reinforced participants learning. Temporal predictions strongly modulate motor preparation (Rolke & Ulrich, 2010; Thomaschke et al., 2011; Volberg & Thomaschke, 2017), and coupling of motor actions to temporally predictable events enhances perceptual modulations (Coull et al., 2024). Future research should confirm whether temporally predictable motor responses are necessary for temporal regularities to be learned and to benefit behaviour, as has been suggested by some researchers (e.g., Salet et al., 2022; Thomaschke & Dreisbach, 2013). Analysing the effects of different combinations of motor, colour, and temporal regularities will be essential for unpacking their individual contributions.

Our task and eye tracking approach offer a promising method for further investigation of feature-temporal attentional guidance in dynamic environments. For example, this framework could be used to understand the influences on behaviour of temporal regularities operating at different levels; from more local temporal intervals between successive stimuli, to regularities occurring across multiple trials.

Future research could also explore different combinations of temporally contingent feature-, location-, and action-based regularities to better understand how they interact to enhance search efficiency. It would be valuable to determine whether, when concurrent, these regularities provide additive benefits, if one is dominant and required, or if certain interactions are particularly strong. Additionally, it would be informative to investigate whether temporal predictability alone can guide dynamic visual search behaviour, without co-occurrence with another sensory-motor prediction.

Finally, future work should examine the paths via which predictable timing can enhance search performance through dynamic attentional guidance. While we focussed on improvements driven by increased attentional capture by temporally expected target features, it remains unclear whether similar benefits could arise from dynamic suppression of temporally likely *distractor* features. This is an intuitive possibility, as dynamic suppression has been demonstrated for salient distractors appearing at predictable times (Lamy, 2005) and at temporally-likely locations (Xu et al., 2021). Investigating suppression mechanisms could provide further insight into how temporal expectations shape attentional control.

## Conclusion

Using a novel dynamic visual-search task, we provide initial evidence for adaptive, temporally-tuned shifts of feature-based attentional guidance during visual search. We found that participants’ learning of temporally contingent regularities affected their task performance, their attentional capture by target features, and their motor response activity in a temporally specific way. This work underscores the flexibility of our attentional system and the utility of temporal predictions in modulating attentional guidance, even under large spatial uncertainty and within crowded dynamic environments.

## Acknowledgments

The authors would like to thank Daniela Gresch and Irene Echeverria Altuna for their help with EMG analysis, other members of the Brain and Cognition Lab for many insightful discussions, and the participants who gave their time to participate in this study. This research was supported by an Oxford Medical Sciences Graduate School Studentship to G.C.W (funded by EPSRC and the Department of Experimental Psychology, University of Oxford); a Wellcome Trust Senior Investigator Award to A.C.N. (104571/Z/14/Z). For the purpose of open access, the author has applied a CC BY public copyright licence to any Author Accepted Manuscript version arising from this submission.

## Supplementary Materials

**Supplementary Figure 1.**
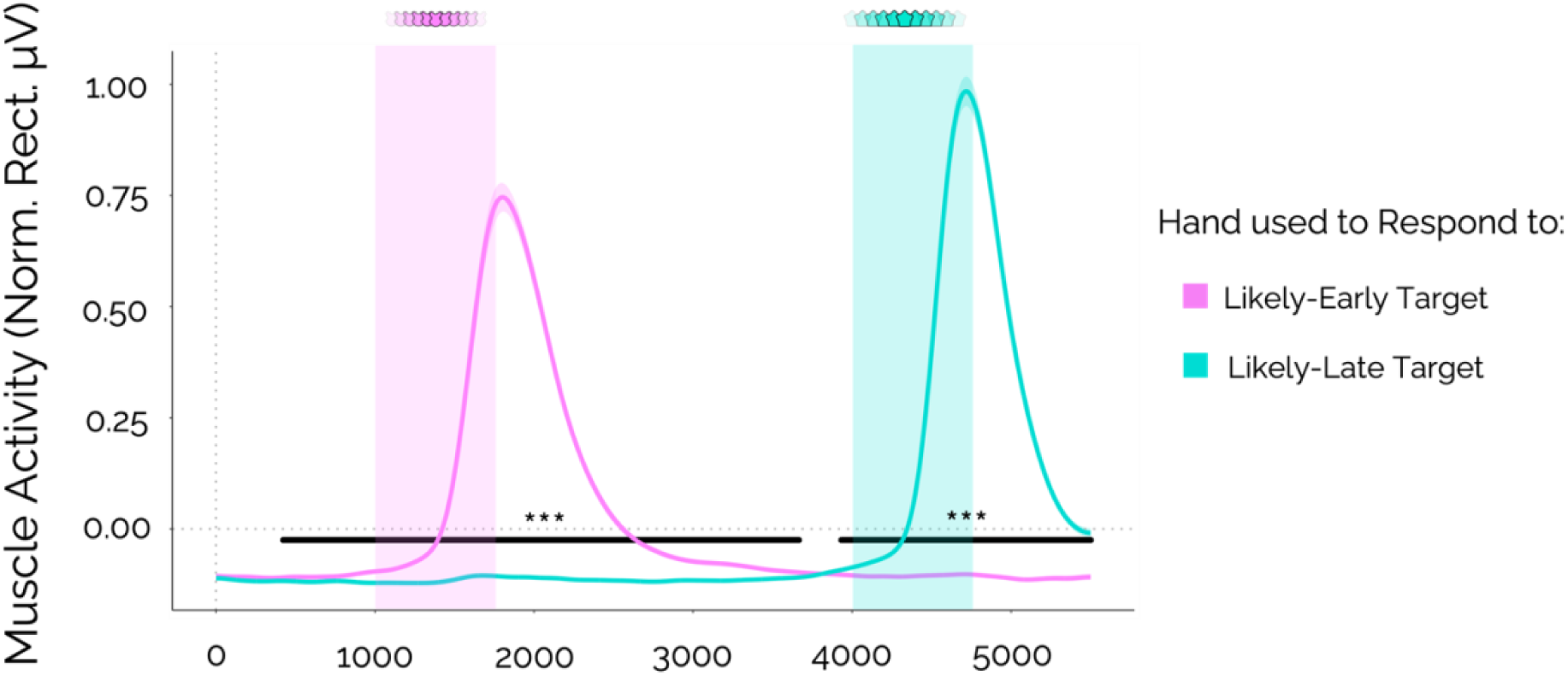
Normalised rectified muscle activity in hands used to respond to the likely-early and likely-late targets over the 5500 ms duration trials where both targets appeared, across all learning blocks, recorded using EMG. The solid pink and blue lines represent the mean values and the shaded areas around them represent ±1 SE. The pink and blue columns indicate the time windows over which the likely-early and likely-late targets, respectively, appeared. The two black horizontal lines mark clusters of significant differences in activity between the two hands, where *** = *p* < .001. For the earlier cluster, the peak difference occurred at 1804 ms. For the later cluster, the peak difference occurred at 4717 ms.

**Supplementary Figure 2.**
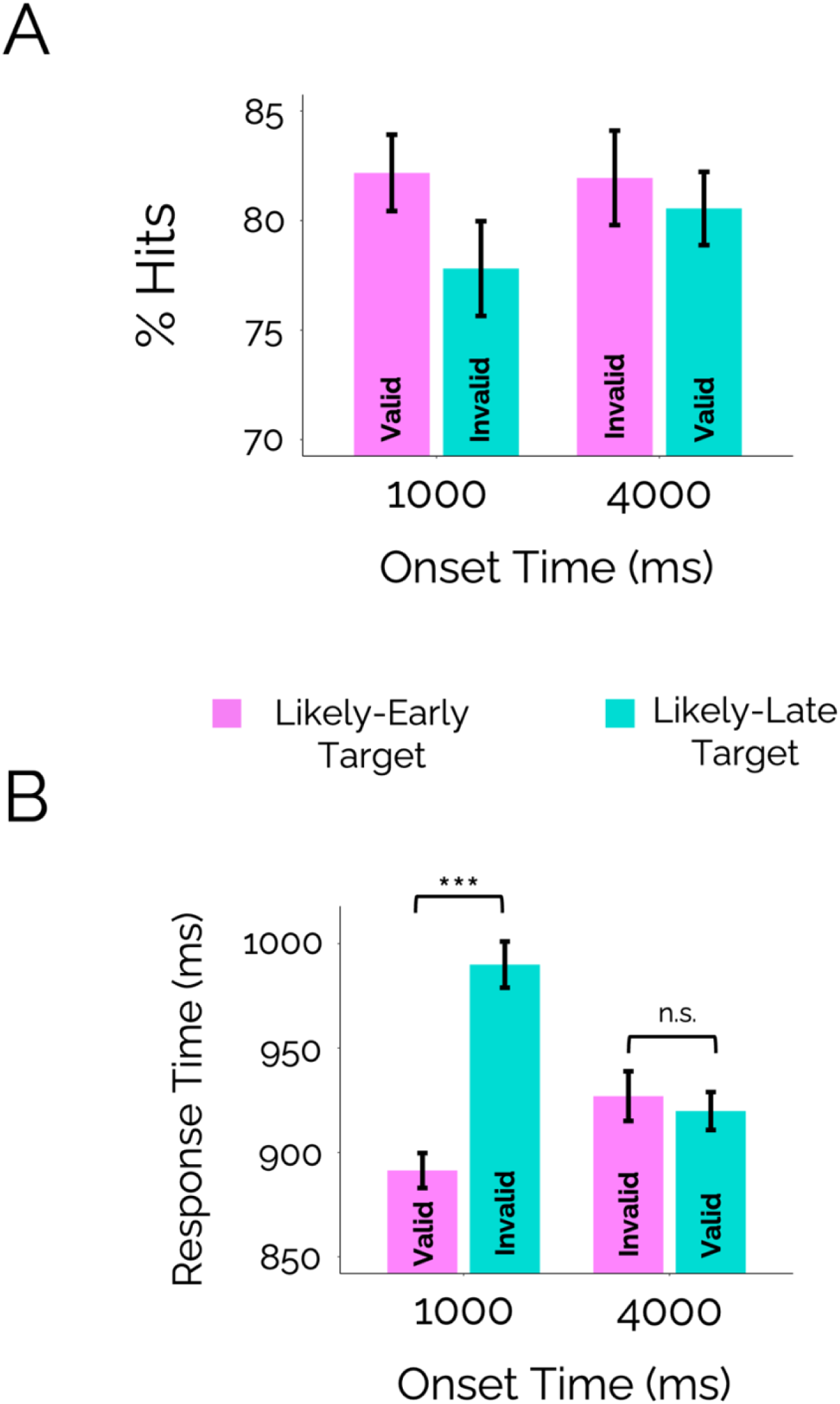
(A) Mean hits for identifying targets in the testing block, for likely-early and likely-late targets appearing at each onset time. Error bars represent ±1 standard error (SE) around the mean. A 2x2 ANOVA analysing the effects of target likelihood and onset time revealed that neither factor, nor their interaction, significantly affected hits. The full output of this ANOVA is shown in Supplementary Table 1 below. (B) Mean response times (RTs) for identifying targets in the testing block, for likely-early and likely-late targets appearing at each onset time. A 2x2 (target likelihood x onset time) ANOVA revealed a significant main effect of target likelihood on RTs (*p* < .001). Overall, participants were significantly faster to identify the likely-early target (M = 897 ms, SD = 49 ms) than the likely-late target (M = 950 ms, SD = 49 ms). The ANOVA also revealed a significant interaction between target likelihood and onset time (*p* < .001). A follow-up t-test revealed that, when targets appeared at 1000 ms, target likelihood significantly affected RTs (*t*(47) = -6.98, *p* < .001, *d* = -1.01, BF_10_ = 1426358.00) whereby the likely-early target (M = 891 ms, SD = 69 ms) was identified significantly faster than the likely-late target (M = 990 ms, SD = 69 ms). A second follow-up t-test revealed that, when targets appeared at 4000 ms, target likelihood had no significant effect on RTs (*t*(47) = 0.45, *p* = .652, *d* = 0.07, BF_10_ = 0.17). The statistical significance of comparisons is indicated in the figure with asterisks: *** = *p* < .001, n.s. = not significant. Full output of the ANOVA analysing RTs is shown in Supplementary Table 2 below.

**Supplementary Table 1.** Hits ANOVA Output.

Hits ANOVA Output
| Predictor | $F$ | $df_1$ | $df_2$ | $p$ | $\eta^2_G$ |
| --- | --- | --- | --- | --- | --- |
| Onset Time | 0.60 | 1 | 47 | .442 | .002 |
| Target Likelihood | 1.40 | 1 | 47 | .243 | .011 |
| Onset Time : Target Likelihood | 0.80 | 1 | 47 | .377 | .003 |

**Supplementary Table 2.** RT ANOVA Output.

RT ANOVA Output
| Predictor | $F$ | $df_1$ | $df_2$ | $p$ | $\eta^2_G$ |
| --- | --- | --- | --- | --- | --- |
| Onset Time | 3.37 | 1 | 47 | .073 | .009 |
| **Target Likelihood** | **17.50** | **1** | **47** | **< .001** | **.063** |
| **Onset Time : Target Likelihood** | **26.77** | **1** | **47** | **< .001** | **.082** |

*Note.* Significant predictors are shown in bold.

**Supplementary Table 3.** Colour Type Custom Contrasts for Distractor Fixations Analysis.

Colour Type Custom Contrasts for Distractor Fixations Analysis
| Colour Type Level | Goal Dependent Contrast<br>(No Colour Match vs.<br>Target Colour Match) | Prediction Contrast<br>(Likely-Early vs. Likely-Late<br>Target Colour Match) |
| --- | --- | --- |
| No Match | 1 | 0 |
| Likely-Early Target Match | -0.5 | -0.5 |
| Likely-Late Target Match | -0.5 | 0.5 |

*Note.* The above contrasts were designed to explore two specific comparisons of the three levels of the colour type variable. The levels code whether a distractor shared its colour with neither target (No Match), the likely-early target (Likely-Early Target Match) or the likely-late target (Likely-Late Target Match). The first comparison, ‘Goal Dependent Contrast (No Colour Match vs. Target Colour Match)’, compared distractors which did not share their colour with either target to those which shared their colour with any target. The second comparison, ‘Prediction Contrast (Likely-Early vs. Likely-Late Target Colour Match)’, compared distractors which shared their colour with the likely-early target to those which shared their colour with the likely-late target.

**Supplementary Figure 3.**
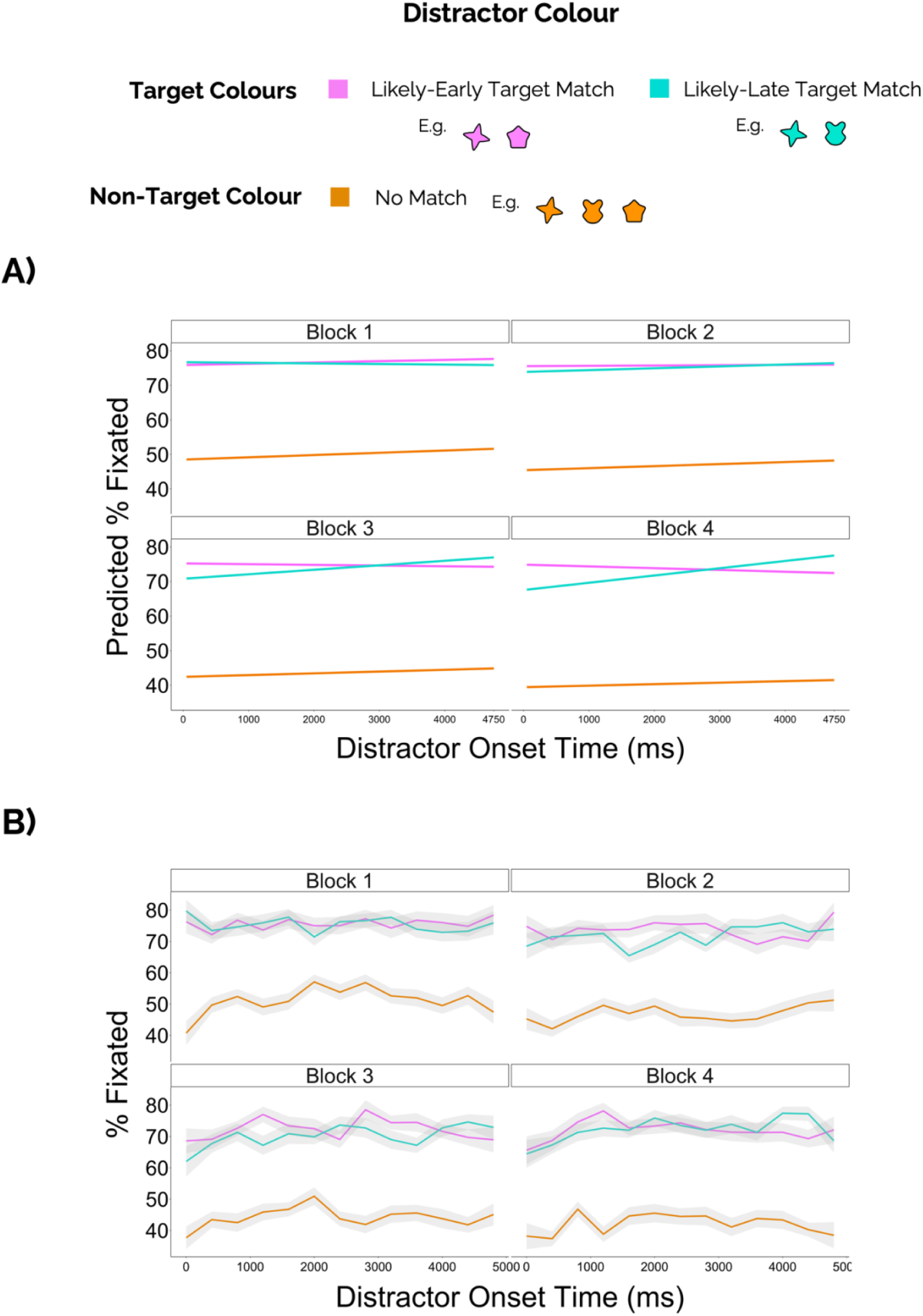
(A) Visualisation of the fixed effects output of the GLMM giving the predicted percentage of distractors fixated during target absent trials across levels of colour type, onset times, and all learning blocks. (B) The actual recorded percentage of distractors fixated across levels of colour type, onset times, and all learning blocks. This plot only includes data from target absent trials of the learning session. The solid pink, blue, and orange lines represent the mean values, while the grey shaded areas represent ±1 SE around the mean. For this plot only, data were smoothed by rounding onset times to the nearest 400 ms.

## References

Albers, C., & Lakens, D. (2018). When power analyses based on pilot data are biased: Inaccurate effect size estimators and follow-up bias. Journal of Experimental Social Psychology, 74, 187–195. 10.1016/J.JESP.2017.09.004

Alexander, R. G., Nahvi, R. J., & Zelinsky, G. J. (2019). Specifying the precision of guiding features for visual search. Journal of Experimental Psychology: Human Perception and Performance, 45(9), 1248–1264. 10.1037/xhp0000668

Anderson, B. A., Kim, H., Kim, A. J., Liao, M.-R., Mrkonja, L., Clement, A., & Grégoire, L. (2021). The past, present, and future of selection history. Neuroscience & Biobehavioral Reviews, 130, 326–350. 10.1016/j.neubiorev.2021.09.004

Awh, E., Belopolsky, A. V., & Theeuwes, J. (2012). Top-down versus bottom-up attentional control: a failed theoretical dichotomy. Trends in Cognitive Sciences, 16(8), 437–443. 10.1016/j.tics.2012.06.010

Bangert, A. S., Kurby, C. A., Hughes, A. S., & Carrasco, O. (2020). Crossing event boundaries changes prospective perceptions of temporal length and proximity. *Attention, Perception*, & Psychophysics, 82(3), 1459–1472. 10.3758/s13414-019-01829-x

Bates, D., Kliegl, R., Vasishth, S., & Baayen, H. (2015). Parsimonious Mixed Models. ArXiv, 1506.

Bates, D., Mächler, M., Bolker, B., & Walker, S. (2015). Fitting Linear Mixed-Effects Models Using **lme4**. Journal of Statistical Software, 67(1). 10.18637/jss.v067.i01

Boettcher, S. E. P., & Nobre, A. C. (2025). Going through the motions: Biasing of dynamic attentional templates. Journal of Experimental Psychology: General, 154(1), 111–127. 10.1037/xge0001665

Boettcher, S. E. P., Shalev, N., Wolfe, J. M., & Nobre, A. C. (2022). Right place, right time: Spatiotemporal predictions guide attention in dynamic visual search. Journal of Experimental Psychology: General, 151(2), 348–362. 10.1037/xge0000901

Brignani, D., Lepsien, J., & Nobre, A. C. (2010). Purely endogenous capture of attention by task-defining features proceeds independently from spatial attention. NeuroImage, 51(2), 859–866. 10.1016/j.neuroimage.2010.03.029

Coull, J. T., Frith, C. D., Büchel, C., & Nobre, A. C. (2000). Orienting attention in time: behavioural and neuroanatomical distinction between exogenous and endogenous shifts. Neuropsychologia, 38(6), 808–819. 10.1016/S0028-3932(99)00132-3

Coull, J. T., Korolczuk, I., & Morillon, B. (2024). The Motor of Time: Coupling Action to Temporally Predictable Events Heightens Perception (pp. 199–213). 10.1007/978-3-031-60183-5_11

Echeverria-Altuna, I., Nobre, A. C., & Boettcher, S. E. P. (2024). Goal-Dependent Use of Temporal Regularities to Orient Attention under Spatial and Action Uncertainty. Journal of Cognition, 7(1), 37. 10.5334/joc.360

Frossard, J., & Renaud, O. (2021). Permutation Tests for Regression, ANOVA, and Comparison of Signals: The **permuco** Package. Journal of Statistical Software, 99(15). 10.18637/jss.v099.i15

Gagné, M., & Schneider, C. (2008). Dynamic influence of wrist flexion and extension on the intracortical inhibition of the first dorsal interosseus muscle during precision grip. Brain Research, 1195, 77–88. 10.1016/j.brainres.2007.12.021

Gramfort, A. (2013). MEG and EEG data analysis with MNE-Python. Frontiers in Neuroscience, 7. 10.3389/fnins.2013.00267

Harrison, X. A., Donaldson, L., Correa-Cano, M. E., Evans, J., Fisher, D. N., Goodwin, C. E. D., Robinson, B. S., Hodgson, D. J., & Inger, R. (2018). A brief introduction to mixed effects modelling and multi-model inference in ecology. PeerJ, 6, e4794. 10.7717/peerj.4794

Heideman, S. G., Rohenkohl, G., Chauvin, J. J., Palmer, C. E., van Ede, F., & Nobre, A. C. (2018). Anticipatory neural dynamics of spatial-temporal orienting of attention in younger and older adults. NeuroImage, 178, 46–56. 10.1016/j.neuroimage.2018.05.002

Hollingworth, A. (2012). Guidance of Visual Search by Memory and Knowledge. *Nebraska Symposium on Motivation*. Nebraska Symposium on Motivation, 59, 63. /pmc/articles/PMC3875155/

Hollingworth, A., & Bahle, B. (2020). Eye tracking in visual search experiments. Neuromethods, 151, 23–35. 10.1007/7657_2019_30/COVER

Inc., T. M. (2018). MATLAB version: 9.4.0 (R2018a). The MathWorks Inc. https://www.mathworks.com

Kleim, J. A., Kleim, E. D., & Cramer, S. C. (2007). Systematic assessment of training-induced changes in corticospinal output to hand using frameless stereotaxic transcranial magnetic stimulation. Nature Protocols, 2(7), 1675–1684. 10.1038/nprot.2007.206

Kliegl. (2010). Experimental effects and individual differences in linear mixed models: estimating the relationship between spatial, object, and attraction effects in visual attention. Frontiers in Psychology. 10.3389/fpsyg.2010.00238

Kumle, L., Võ, M. L., & Draschkow, D. (2021). Estimating power in (generalized) linear mixed models: an open introduction and tutorial in R. B. Behavior Research Methods. 10.3758/s13428-021-01546-0

Kwon, S., Bahn, S., Ahn, S. H., Lee, Y., & Yun, M. H. (2016). A study on the relationships among hand muscles and form factors of large-screen curved mobile devices. International Journal of Industrial Ergonomics, 56, 17–24. 10.1016/j.ergon.2016.07.003

Lamy, D. (2005). Temporal expectations modulate attentional capture. Psychonomic Bulletin & Review, 12(6), 1112–1119. 10.3758/BF03206452

Lawrence, M. A. (2016). *ez: Easy Analysis and Visualization of Factorial Experiments*. https://CRAN.R-project.org/package=ez

Li, A. Y., Liang, J. C., Lee, A. C. H., & Barense, M. D. (2020). The validated circular shape space: Quantifying the visual similarity of shape. Journal of Experimental Psychology: General, 149(5), 949–966. 10.1037/xge0000693

Morey, R. D., & Rouder, J. N. (2022). *BayesFactor: Computation of Bayes Factors for Common Designs*. https://CRAN.R-project.org/package=BayesFactor

Muhl-Richardson, A., Cornes, K., Godwin, H. J., Garner, M., Hadwin, J. A., Liversedge, S. P., & Donnelly, N. (2018). Searching for two categories of target in dynamic visual displays impairs monitoring ability. Applied Cognitive Psychology, 32(4), 440–449. 10.1002/ACP.3416

Muhl-Richardson, A., Godwin, H. J., Garner, M., Hadwin, J. A., Liversedge, S. P., & Donnelly, N. (2018). Individual differences in search and monitoring for color targets in dynamic visual displays. Journal of Experimental Psychology: Applied, 24(4), 564–577. 10.1037/xap0000155

Ongchoco, J. D. K., Yates, T. S., & Scholl, B. J. (2023). Event segmentation structures temporal experience: Simultaneous dilation and contraction in rhythmic reproductions. Journal of Experimental Psychology: General, 152(11), 3266–3276. 10.1037/xge0001447

Pfeuffer, C. U., Aufschnaiter, S., Thomaschke, R., & Kiesel, A. (2020). Only time will tell the future: Anticipatory saccades reveal the temporal dynamics of time-based location and task expectancy. Journal of Experimental Psychology: Human Perception and Performance, 46(10), 1183–1200. 10.1037/xhp0000850

R Core Team. (2024). *R: A Language and Environment for Statistical Computing*. https://www.R-project.org/

Rolke, B., & Ulrich, R. (2010). *On the locus of temporal preparation: Enhancement of premotor processes*. https://philpapers.org/rec/ROLOTL

Saenz, M., Buracas, G. T., & Boynton, G. M. (2002). Global effects of feature-based attention in human visual cortex. Nature Neuroscience, 5(7), 631–632. 10.1038/nn876

Salet, J. M., Kruijne, W., Rijn, H. van, Zimmermann, E., & Schlichting, N. (2022). Statistical Learning Emerges from Temporally Preparing What Action to Perform Where. BioRxiv, 2022.08.06.502990. 10.1101/2022.08.06.502990

Schreij, D., Owens, C., & Theeuwes, J. (2008). Abrupt onsets capture attention independent of top-down control settings. Perception & Psychophysics, 70(2), 208–218. 10.3758/PP.70.2.208

Schreij, D., Theeuwes, J., & Olivers, C. N. L. (2010). Abrupt onsets capture attention independent of top-down control settings II: Additivity is no evidence for filtering. *Attention, Perception*, & Psychophysics, 72(3), 672–682. 10.3758/APP.72.3.672

Shalev, N., Boettcher, S., & Nobre, A. C. (2024). Spatiotemporal predictions guide attention throughout the adult lifespan. Npj Science of Learning, 9(1), 70. 10.1038/s41539-024-00281-3

Shalev, N., Boettcher, S., Wilkinson, H., Scerif, G., & Nobre, A. C. (2022). Be there on time: Spatial-temporal regularities guide young children’s attention in dynamic environments. Child Development, 93(5), 1414–1426. 10.1111/cdev.13770

Thomaschke, R., & Dreisbach, G. (2013). Temporal predictability facilitates action, not perception. Psychological Science, 24(7), 1335–1340. 10.1177/0956797612469411

Thomaschke, R., Hoffmann, J., Haering, C., & Kiesel, A. (2016). Time-Based Expectancy for Task Relevant Stimulus Features. Timing & Time Perception, 4(3), 248–270. 10.1163/22134468-00002069

Thomaschke, R., Kiesel, A., & Hoffmann, J. (2011). Response specific temporal expectancy: evidence from a variable foreperiod paradigm. Attention, Perception & Psychophysics, 73(7), 2309–2322. 10.3758/S13414-011-0179-6

Townsend, J. T., & Ashby, F. G. (1983). Stochastic Modeling of Elementary Psychological Processes. Cambridge University Press.

Treue, S., & Trujillo, J. C. M. (1999). Feature-based attention influences motion processing gain in macaque visual cortex. Nature, 399(6736), 575–579. 10.1038/21176

Virtanen, P., Gommers, R., Oliphant, T. E., Haberland, M., Reddy, T., Cournapeau, D., Burovski, E., Peterson, P., Weckesser, W., Bright, J., van der Walt, S. J., Brett, M., Wilson, J., Millman, K. J., Mayorov, N., Nelson, A. R. J., Jones, E., Kern, R., Larson, E., … Vázquez-Baeza, Y. (2020). SciPy 1.0: fundamental algorithms for scientific computing in Python. Nature Methods, 17(3), 261–272. 10.1038/s41592-019-0686-2

Volberg, G., & Thomaschke, R. (2017). Time-based expectations entail preparatory motor activity. Cortex, 92, 261–270. 10.1016/J.CORTEX.2017.04.019

Wagener, A., & Hoffmann, J. (2010). Temporal Cueing of Target-Identity and Target-Location. 10.1027/1618-3169/A000054, 57(6), 436–445. 10.1027/1618-3169/A000054

Wickham, H. (2016). ggplot2: Elegant Graphics for Data Analysis. Springer-Verlag New York. https://ggplot2.tidyverse.org

Wolfe, J. M. (2020). Annual Review of Vision Science Visual Search: How Do We Find What We Are Looking For? 10.1146/annurev-vision-091718

Wolfe, J. M. (2021). Guided Search 6.0: An updated model of visual search. Psychonomic Bulletin & Review 2021 28:4, 28(4), 1060–1092. 10.3758/S13423-020-01859-9

Wolfe, J. M., & Horowitz, T. S. (2017). Five factors that guide attention in visual search.Nature Human Behaviour 2017 1:3, 1(3), 1–8. 10.1038/s41562-017-0058

Xu, Z., Los, S. A., & Theeuwes, J. (2021). Attentional suppression in time and space. Journal of Experimental Psychology: Human Perception and Performance, 47(8), 1056–1062. 10.1037/XHP0000925

Xu, Z., Theeuwes, J., & Los, S. A. (2023). Statistical learning of spatiotemporal regularities dynamically guides visual attention across space. *Attention, Perception*, & Psychophysics, 85(4), 1054–1072. 10.3758/s13414-022-02573-5

Xu, Z., Theeuwes, J., & Los, S. A. (2025). Statistical learning of spatiotemporal target regularities in the absence of saliency. *Attention, Perception*, & Psychophysics. 10.3758/s13414-024-02992-6

